## Supplementary Data for "Mapping patterns of thought onto brain activity during movie-watching" (from another version of this preprint: version 3, 2024-08-06)

### Supplementary Tables

### Supplementary Table 1

*Multi-dimensional Experience sampling (mDES)*

| Questions | Label | Low | High |
|---|---|---|---|
| My thoughts were focused on an external task or activity: | Focus | Not at all | Completely |
| My thoughts involved future events: | Future | Not at all | Completely |
| My thoughts involved past events: | Past | Not at all | Completely |
| My thoughts involved myself: | Self | Not at all | Completely |
| My thoughts involved other people: | Other | Not at all | Completely |
| The emotion of my thoughts was: | Emotion | Negative | Positive |
| My thoughts involved images | Images | Not at all | Completely |
| My thoughts were detailed and specific: | Detailed | Not at all | Completely |
| My thoughts were: | Deliberate | Spontaneous | Deliberate |
| I was thinking about solutions to problems (or goals): | Problem | Not at all | Completely |
| My thoughts were intrusive: | Intrusive | Not at all | Completely |
| My thoughts contained information I already knew (e.g., knowledge or memories): | Knowledge | Not at all | Completely |
| I was absorbed in the contents of my thoughts | Absorption | Not at all | Completely |

| | | | |
|---|---|---|---|
| My thoughts were distracting me from what I am doing | Distracting | Not at all | Completely |
| My thoughts involved words | Words | Not at all | Completely |
| My thoughts involved sounds | Sounds | Not at all | Completely |

**Supplementary Table 2**

Percent variance explained by principal components in each movie

| **Movie** | PC1 | PC2 | PC3 | PC4 | Total |
|---|---|---|---|---|---|
| ***Citizenfour*** | 14.01% | 14.97% | 12.85% | 8.51% | 50.34% |
| ***Little Miss Sunshine*** | 14.93% | 13.02% | 13.67% | 9.77% | 51.43% |
| ***500 Days of Summer*** | 18.22% | 11.26% | 12.28% | 9.61% | 51.38% |
| **Combined Dataset** | 26.07% | 10.45% | 7.73% | 6.77% | 51.03% |

**Supplementary Table 3**

*Linear Mixed Models of Variance in Thoughts across Movies*

| | | | | | | |
|---|---|---|---|---|---|---|
| Model 1 | | | | | | |
| | Sum Sq. | Mean Sq. | Num DF | Den DF | F value | p-value |
| Movie | 16.83 | 8.41 | 2 | 2015.30 | 5.41 | 0.005 |
| Model 1 lsmeans | | | | | | |
| | Contrasts | lsmean | SE | df | t-value | p-value |
| | c4 - lms | -0.02 | .10 | 159 | 1.31 | .392 |
| | c4 - summer | -0.11 | .10 | 159 | -1.97 | .121 |
| | lms - summer | 0.11 | .10 | 160 | -3.27 | .003 |
| Model 2 | | | | | | |
| | Sum Sq. | Mean Sq. | Num DF | Den DF | F value | p-value |
| Movie | 186.92 | 93.46 | 2 | 2015.30 | 77.84 | < .001 |
| Model 2 lsmeans | | | | | | |
| | Contrasts | lsmean | SE | df | t-value | p-value |
| | c4 - lms | 0.56 | .06 | 2015 | 9.66 | < .001 |
| | c4 - summer | 0.68 | .06 | 2016 | 11.66 | < .001 |
| | lms - summer | 0.12 | .06 | 2016 | 2.03 | .106 |
| Model 3 | | | | | | |
| | Sum Sq. | Mean Sq. | Num DF | Den DF | F value | p-value |
| Movie | 34.29 | 17.15 | 2 | 2015.40 | 12.90 | < .001 |
| Model 3 lsmeans | | | | | | |
| | Contrasts | lsmean | SE | df | t-value | p-value |
| | c4 - lms | 0.30 | .06 | 2015 | 5.14 | < .001 |
| | c4 - summer | 0.21 | .06 | 2016 | 2.58 | .001 |
| | lms - summer | -0.09 | .06 | 2016 | -1.54 | .271 |
| Model 4 | | | | | | |
| | Sum Sq. | Mean Sq. | Num DF | Den DF | F value | p-value |
| Movie | 149.90 | 74.95 | 2 | 2015.70 | 82.69 | < .001 |
| Model 4 lsmeans | | | | | | |
| | Contrasts | lsmean | SE | df | t-value | p-value |
| | c4 - lms | -0.31 | .05 | 2015 | -6.07 | < .001 |
| | c4 - summer | -0.65 | .05 | 2016 | -12.85 | < .001 |
| | lms - summer | -0.34 | .05 | 2016 | -6.81 | < .001 |

**Supplementary Table 4**

*Movie Comprehension Questions*

| Movie | Question | Answer |
|---|---|---|
| *Citizenfour* | What animal is seen on screen in the first scene? | Dog |
| *Citizenfour* | What president is being discussed in the phone call with David? | Barack Obama |
| *Citizenfour* | After 4 years in the military, who directly entered the NSA with 37 years of combined service? | William |
| *Citizenfour* | According to the movie, what historical event triggered the government to begin spying on citizens? | 11-Sep |
| *Little Miss Sunshine* | What breakfast item did Olive order a la mode? | Waffles |
| *Little Miss Sunshine* | What colour is the Volkswagen Transporter that the family drives? | Yellow |
| *Little Miss Sunshine* | How does the family get into their car after taking it to the repair shop? | Pushing it |
| *Little Miss Sunshine* | What type of language does Richard claim is “the refuge of losers”? | Sarcasm |
| *500 Days of Summer* | What event are both Summer and Tom attending? | Wedding |
| *500 Days of Summer* | What piece of jewelry is Summer showing her friend when Tom abruptly leaves her party? | Engagement ring |
| *500 Days of Summer* | On day 440, what time does Tom’s alarm clock go off? | 7:00 AM |
| *500 Days of Summer* | What book is Tom reading on the train, which he also gifts to Summer at her party? | Architecture of happiness |

**Supplementary Table 5**

*Linear Mixed Models of Comprehension model*

| | Sum Sq. | Mean Sq. | Num Df | Den Df | F-value | p-value |
|---|---|---|---|---|---|---|
| Movie | 47.32 | 23.66 | 2 | 254.12 | 49.33 | < .001 |
| PCA_1 | 0.64 | 0.64 | 1 | 338.84 | 1.32 | .376 |
| PCA_2 | 4.45 | 4.45 | 1 | 324.31 | 8.27 | .011 |
| PCA_3 | 0.01 | 0.01 | 1 | 332.52 | 0.03 | .865 |
| PCA_4 | 3.98 | 3.98 | 1 | 341.44 | 8.30 | .013 |
| Movie*PCA_1 | 4.28 | 2.14 | 2 | 268.96 | 3.36 | .013 |
| Movie*PCA_2 | 1.18 | 0.89 | 2 | 259.95 | 1.85 | .286 |
| Movie*PCA_3 | 1.01 | 0.51 | 2 | 264.49 | 1.05 | .450 |
| Movie*PCA_4 | 0.32 | 0.16 | 2 | 269.18 | 0.33 | .810 |

*Pairwise Comparisons for significant main effect of movie*

| Contrasts | Estimate | SE | Df | t-value | p-value |
|---|---|---|---|---|---|
| c4 - lms | -0.93 | .10 | 249 | -9.16 | < .001 |
| c4 - summer | -0.91 | .11 | 273 | -8.33 | < .001 |
| lms - summer | 0.02 | .10 | 242 | 0.18 | .982 |

*Note.* The abbreviation ‘c4’ refers to *Citizenfour*, ‘lms’ refers to *Little Miss Sunshine*, and ‘summer’ refers to *500 Days of Summer*.

### Supplementary Table 6

*FSL FEAT Query Parameter Estimates*

**All Movies**

| | Mean | LCI | UCI |
|---|---|---|---|
| Episodic Knowledge | 0.62 | 0.27 | 0.97 |
| Intrusive Distraction | -0.78 | -1.37 | -0.20 |
| Verbal Detail | -1.64 | -2.11 | -1.18 |
| Sensory Engagement | 1.26 | 0.81 | 1.71 |

***500 Days of Summer***

| | Mean | LCI | UCI |
|---|---|---|---|
| Episodic Knowledge | -0.23 | -0.58 | 0.12 |
| Intrusive Distraction | -2.34 | -2.89 | -1.80 |
| Verbal Detail | -0.87 | -1.25 | -0.48 |
| Sensory Engagement | 0.18 | -0.02 | 0.39 |

***Citizenfour***

| | Mean | LCI | UCI |
|---|---|---|---|
| Episodic Knowledge | 1.49 | 1.06 | 1.91 |
| Intrusive Distraction | 0.26 | -0.26 | 0.78 |
| Verbal Detail | -2.98 | -3.64 | -2.31 |
| Sensory Engagement | 2.77 | 2.27 | 3.26 |

***Little Miss Sunshine***

| | Mean | LCI | UCI |
|---|---|---|---|
| Episodic Knowledge | 0.88 | -0.15 | 1.90 |
| Intrusive Distraction | 1.29 | -0.75 | 3.33 |
| Verbal Detail | -0.25 | -1.00 | 0.51 |
| Sensory Engagement | 0.29 | -0.60 | 1.19 |

**Supplementary Table 7**

*Grand average of Gradient score by movie*

| | Gradient 1 mean | Gradient 2 mean | Gradient 3 mean | Gradient 4 mean | Gradient 5 mean |
|---|---|---|---|---|---|
| *500 Days of Summer* | -0.00100 | 0.00264 | 0.00201 | 0.00134 | -0.00112 |
| *Citizenfour* | 0.00301 | -0.00191 | 0.00097 | 0.00160 | 0.00000 |
| *Little Miss Sunshine* | 0.00332 | 0.00138 | -0.00099 | 0.00057 | 0.00130 |

**Supplementary Table 8**

*Functional Connectivity Cluster Analysis*

Episodic Knowledge Functional Connectivity

| Cluster Index | Voxels | *p* | log *p* | Z-MAX | Z-MAX X (mm) | Z-MAX Y (mm) | Z-MAX Z (mm) | Z-COG X (mm) | Z-COG Y (mm) | Z-COG Z (mm) |
|---|---|---|---|---|---|---|---|---|---|---|
| 3 | 286 | 3.04E-06 | 5.52 | 4.47 | -8 | -100 | 14 | -8.53 | -97.3 | 13.4 |
| 2 | 184 | 0.000197 | 3.71 | 3.93 | 6 | -92 | 14 | 11.3 | -93.5 | 17.5 |
| 1 | 101 | 0.0109 | 1.96 | 3.84 | -58 | -16 | 6 | -58.2 | -15.2 | 6.22 |

Intrusive Distraction Functional Connectivity

| Cluster Index | Voxels | *p* | log *p* | Z-MAX | Z-MAX X (mm) | Z-MAX Y (mm) | Z-MAX Z (mm) | Z-COG X (mm) | Z-COG Y (mm) | Z-COG Z (mm) |
|---|---|---|---|---|---|---|---|---|---|---|
| 4 | 547 | 1.11E-09 | 8.95 | 4.8 | -34 | -58 | 50 | -36 | -52.6 | 46.4 |
| 3 | 408 | 5.96E-08 | 7.22 | 4.3 | -46 | 8 | 26 | -45.9 | 6.9 | 29.3 |
| 2 | 289 | 4.89E-06 | 5.31 | 4.04 | 42 | -46 | 44 | 38.4 | -51.3 | 45.6 |
| 1 | 130 | 0.00338 | 2.47 | 4.08 | 46 | 48 | -4 | 45 | 48.7 | 1.19 |

Verbal Detail Functional Connectivity

| Cluster Index | Voxels | *p* | log *p* | Z-MAX | Z-MAX X (mm) | Z-MAX Y (mm) | Z-MAX Z (mm) | Z-COG X (mm) | Z-COG Y (mm) | Z-COG Z (mm) |
|---|---|---|---|---|---|---|---|---|---|---|
| 4 | 1895 | 7.05E-24 | 23.2 | 5.24 | -60 | -16 | 6 | -56.3 | -23.3 | 4.47 |
| 3 | 1010 | 2.10E-15 | 14.7 | 4.74 | 58 | -8 | 4 | 58.3 | -18.4 | 3.45 |
| 2 | 105 | 0.00967 | 2.01 | 3.82 | 12 | -88 | 10 | 10.6 | -86.9 | 11 |
| 1 | 75 | 0.05 | 1.3 | 3.74 | -52 | -76 | 6 | -50.3 | -76.6 | 4.32 |

Sensory Engagement Functional Connectivity

| Cluster Index | Voxels | *p* | log *p* | Z-MAX | Z-MAX X (mm) | Z-MAX Y (mm) | Z-MAX Z (mm) | Z-COG X (mm) | Z-COG Y (mm) | Z-COG Z (mm) |
|---|---|---|---|---|---|---|---|---|---|---|
| 5 | 4566 | 7.43E-44 | 43.1 | 5.23 | 6 | -92 | 16 | 9.59 | -80.2 | 1.36 |
| 4 | 340 | 4.77E-07 | 6.32 | 4.22 | -46 | -80 | 2 | -46.4 | -76.7 | 5.65 |
| 3 | 330 | 6.56E-07 | 6.18 | 4.46 | 62 | -8 | 4 | 61.3 | -10.7 | 7.1 |
| 2 | 290 | 2.92E-06 | 5.53 | 4.59 | -64 | -12 | 6 | -61.1 | -13.4 | 5.4 |
| 1 | 156 | 0.000754 | 3.12 | 3.68 | -8 | -60 | 66 | -6.45 | -56.5 | 58.4 |

**Supplementary Table 9**

*Neurosynth Decoder Analysis*

| Episodic Knowledge | | Intrusive Distraction | | Verbal Detail | | Sensory Engagement | |
|---|---|---|---|---|---|---|---|
| association term | *r* | association term | *r* | association term | *r* | association term | *r* |
| cuneus | 0.17 | working | 0.33 | auditory cortex | 0.65 | v1 | 0.40 |
| v1 | 0.17 | working memory | 0.33 | sounds | 0.65 | visual | 0.35 |
| primary visual | 0.13 | intraparietal | 0.31 | auditory | 0.64 | visual cortex | 0.34 |
| sighted | 0.11 | intraparietal sulcus | 0.30 | primary auditory | 0.62 | primary visual | 0.32 |
| pointing | 0.08 | parietal cortex | 0.30 | heschl | 0.61 | occipital | 0.27 |
| visual | 0.08 | tasks | 0.29 | sound | 0.61 | extrastriate | 0.25 |
| occipital | 0.07 | task | 0.27 | heschl gyrus | 0.60 | early visual | 0.25 |
| videos | 0.07 | posterior parietal | 0.27 | speech | 0.60 | mt | 0.24 |
| occipital gyrus | 0.06 | sulcus ips | 0.25 | listening | 0.59 | sighted | 0.23 |
| visual stream | 0.05 | parietal | 0.25 | pitch | 0.59 | motion | 0.21 |
| prefrontal | -0.02 | resting state | -0.18 | prefrontal | -0.16 | prefrontal | -0.08 |
| cerebellum | -0.02 | resting | -0.18 | prefrontal cortex | -0.13 | cingulate | -0.08 |
| premotor | -0.02 | amygdala | -0.11 | cingulate | -0.06 | prefrontal cortex | -0.07 |
| memory | -0.02 | controls | -0.11 | medial | -0.05 | cerebellum | -0.05 |
| movements | -0.02 | emotional | -0.11 | parietal | -0.04 | motor | -0.05 |
| motor | -0.02 | disorder | -0.10 | memory | -0.04 | sensorimotor | -0.05 |
| primary motor | -0.02 | orbitofrontal | -0.08 | cerebellum | -0.04 | primary motor | -0.05 |

| | | | | | | | |
|---|---|---|---|---|---|---|---|
| movement | -0.02 | stimulation | -0.06 | cerebellar | -0.04 | memory | -0.05 |
| parietal | -0.02 | cingulate | -0.06 | disorder | -0.04 | movement | -0.05 |
| pain | -0.02 | disorders | -0.05 | task | -0.03 | motor cortex | -0.05 |

**Supplementary Table 10**

*Linear Mixed Models of Gradients 1-5 Fixed Effects for each Thought Pattern*

**DV: Episodic Knowledge**

| | Estimate | Std. Error | Df | t-value | p-value |
|---|---|---|---|---|---|
| (Intercept) | 0.01 | 0.09 | 118.6 | 0.07 | .940 |
| gradient1 | -0.01 | 0.03 | 2043 | -0.26 | .796 |
| gradient2 | -0.04 | 0.04 | 2043 | -0.94 | .579 |
| gradient3 | -0.03 | 0.04 | 2042 | -0.69 | .613 |
| gradient4 | 0.18 | 0.06 | 2046 | 3.02 | .013 |
| gradient5 | -0.18 | 0.08 | 2045 | -2.24 | .063 |

**DV: Intrusive Distraction**

| | Estimate | Std. Error | Df | t-value | p-value |
|---|---|---|---|---|---|
| (Intercept) | -0.01 | 0.08 | 118.8 | -0.08 | .934 |
| gradient1 | 0.02 | 0.03 | 2045 | 0.68 | .500 |
| gradient2 | 0.05 | 0.04 | 2045 | 1.34 | .412 |
| gradient3 | -0.04 | 0.03 | 2043 | -1.16 | .412 |
| gradient4 | 0.12 | 0.05 | 2048 | 2.17 | .152 |
| gradient5 | -0.06 | 0.07 | 2046 | -0.89 | .470 |

**DV: Verbal Detail**

| | Estimate | Std. Error | Df | t-value | p-value |
|---|---|---|---|---|---|
| (Intercept) | 0.01 | 0.09 | 119 | 0.05 | .963 |
| gradient1 | -0.01 | 0.03 | 2043 | -0.27 | .787 |
| gradient2 | -0.04 | 0.04 | 2043 | -1.03 | .382 |
| gradient3 | 0.03 | 0.03 | 2042 | 1.03 | .382 |
| gradient4 | 0.09 | 0.05 | 2046 | 1.71 | .217 |
| gradient5 | -0.15 | 0.07 | 2045 | -2.06 | .199 |

**DV: Sensory Engagement**

| | Estimate | Std. Error | Df | t-value | p-value |
|---|---|---|---|---|---|
| (Intercept) | 0.01 | 0.07 | 119.15 | 0.16 | .870 |
| gradient1 | -0.09 | 0.03 | 2046 | -3.26 | .006 |
| gradient2 | 0.03 | 0.03 | 2046 | 0.78 | .432 |
| gradient3 | 0.03 | 0.03 | 2044 | 1.17 | .404 |
| gradient4 | 0.07 | 0.05 | 2049 | 1.55 | .204 |
| gradient5 | 0.05 | 0.06 | 2048 | 0.86 | .432 |

**Supplementary Figures**

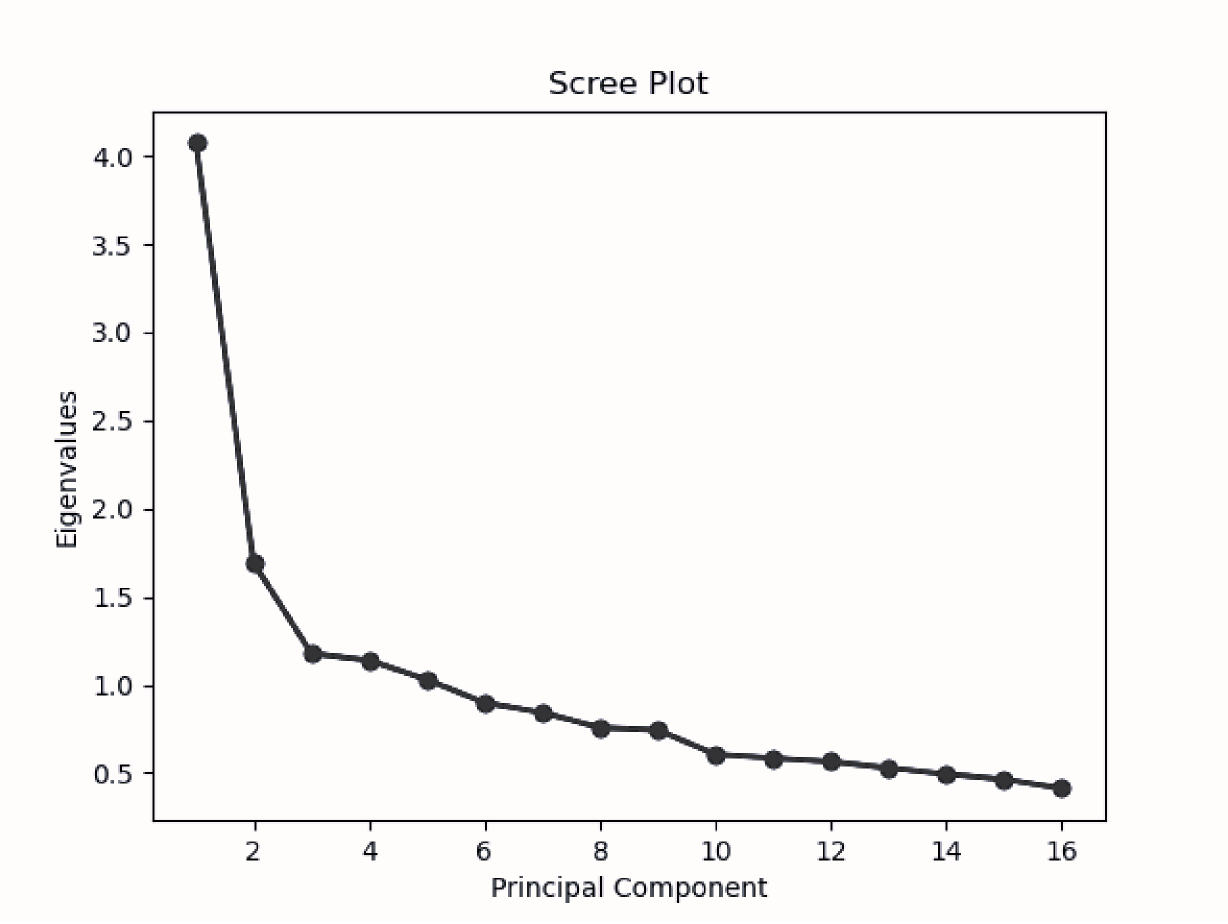

**Supplementary Figure 1.**

The scree plot illustrates the eigenvalues of principal components obtained from a PCA analysis. Each point on the plot represents a principal component, and the y-axis displays the corresponding eigenvalues. The eigen value threshold of 1 and the "elbow" of the plot, where eigenvalues start to level off, is indicative of the optimal number of components to retain. In this case, the analysis suggests a 4-component solution. The scree plot aids in determining the most meaningful components for capturing variance in the data, facilitating a more parsimonious representation of the underlying structure.

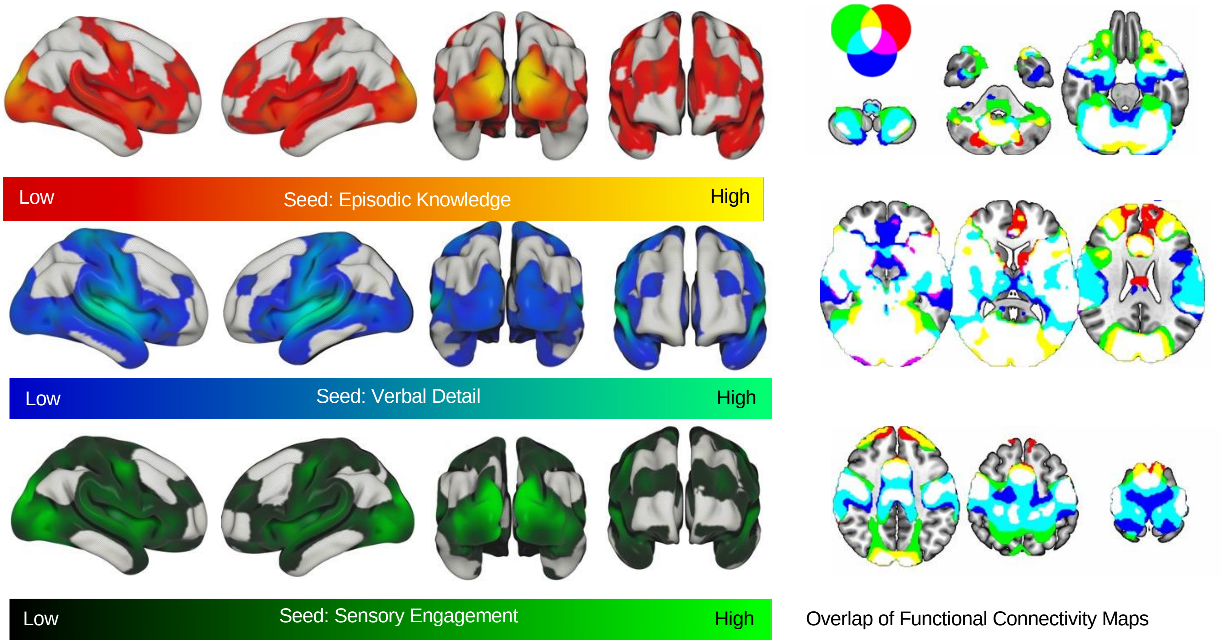

**Supplementary Figure 2.**

Results of functional connectivity (FC) in which we used the regions identified in our voxel space analysis that related to "Episodic Knowledge" (Red - Yellow color map), "Verbal Detail" (Blue-Green color map) and "Sensory Engagement" (Black – Green color map). These are thresholded at $p < .001$ and corrected for FEW at $p < .05$.The panel on the right shows the same fully saturated maps to highlight their common regions (shown in white).

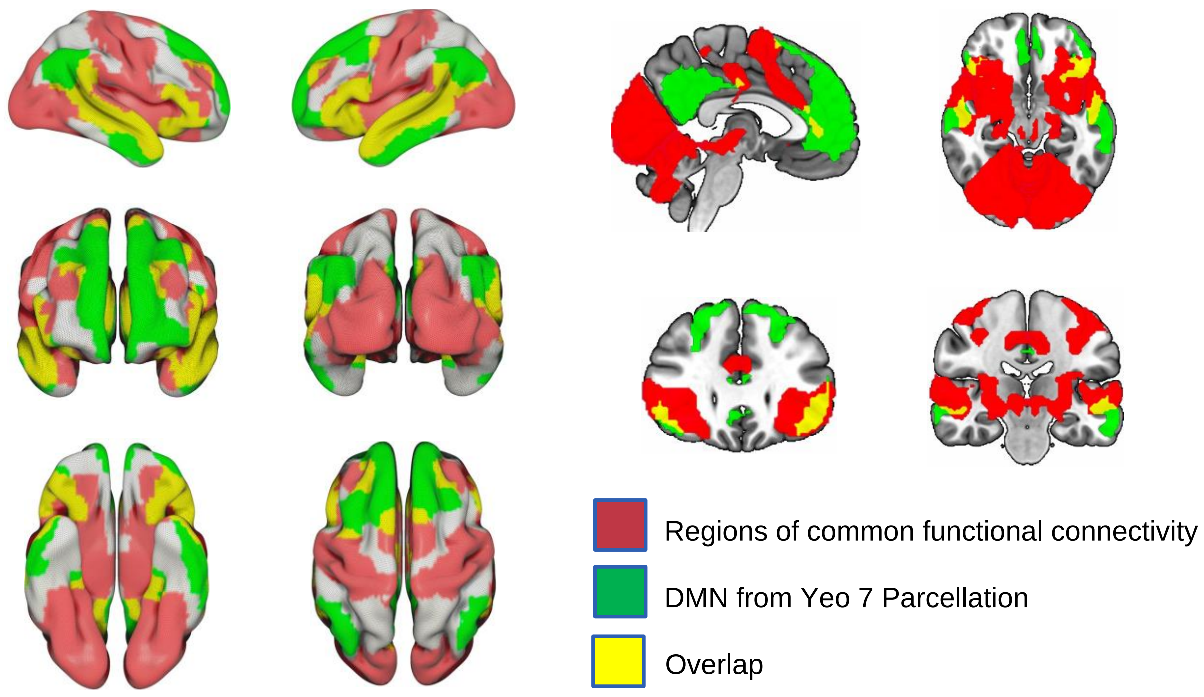

**Supplementary Figure 3.**

Overlap between the regions showing common functional connectivity between regions associated with three mDES components (“Episodic Knowledge”, “Verbal Detail” & “Sensory Engagement”) in the voxel space analysis and the DMN as defined by [45]. Regions in red show regions of common functional connectivity, regions in green are the DMN and regions in yellow are the overlap between these two maps.

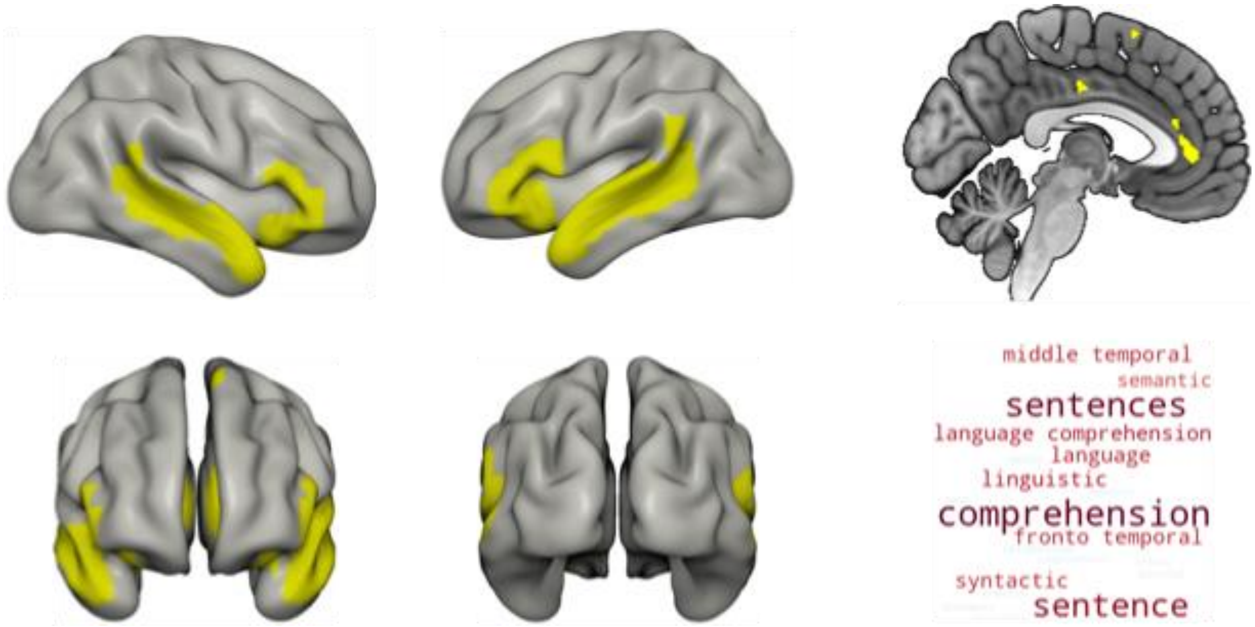

**Supplementary Figure 4**

Meta analytic decoding of the regions showing common functional connectivity between regions associated with three mDES components ("Episodic Knowledge", "Verbal Detail" & "Sensory Engagement") in the voxel space analysis and the DMN as defined by [45] (shown in yellow). The word cloud on the right shows the functional terms most likely to be associated with these regions following a Neurosynth analytic decoding. Regions in red are more likely and regions in blue are less likely. Words in a larger font size are more important.

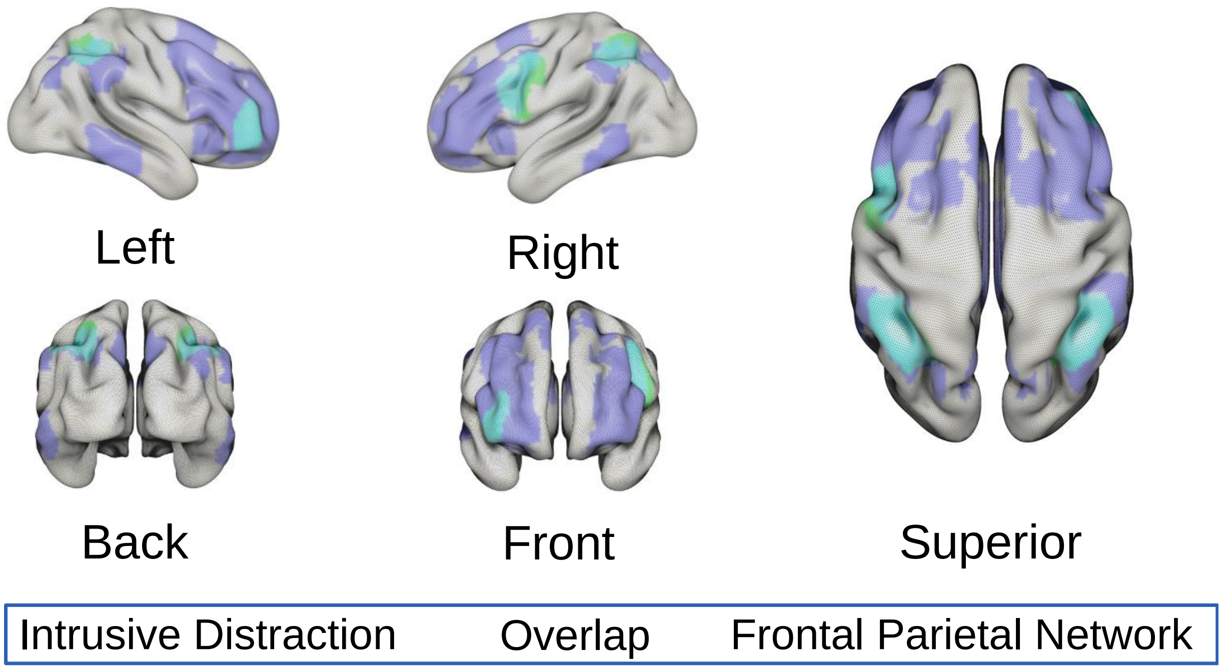

**Supplementary Figure 5**

Overlap between regions associated with “Intrusive Distraction” in our voxel space analysis (green) and the FPN as defined by [45] (lavender). Regions of overlap are presented in cyan.

| Thought Component | Coronal View | Sagittal View | Axial View |
| --- | --- | --- | --- |
| Episodic Knowledge | 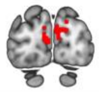 | 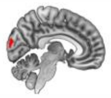 | 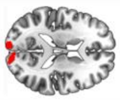 |
| Intrusive Distraction | 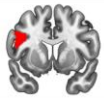 | 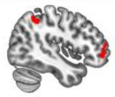 | 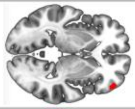 |
| Verbal Detail | 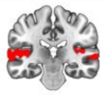 | 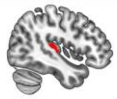 | 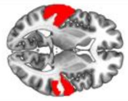 |
| Sensory Engagement | 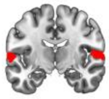 | 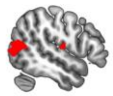 | 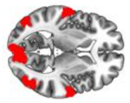 |

**Supplementary Figure 6**

Brain maps of each voxel-space analysis output for each of the corresponding thought components separately, “Episodic Knowledge,” “Intrusive Distraction,” “Verbal Detail,” and “Sensory Engagement.” Each brain map is displayed from the coronal view, sagittal view, and axial view to maximize the understanding of the spatial distribution and localization of related brain regions for each thought pattern.

| Time | Probe 1 | Probe 2 | Probe 3 | Probe 4 | Probe 5 |
|---|---|---|---|---|---|
| **Order 1** | 1:15 | 3:45 | 5:30 | 7:15 | 9:30 |
| **Order 2** | 2:45 | 4:30 | 6:00 | 8:15 | 9:45 |
| **Order 3** | 1:30 | 3:15 | 5:15 | 7:30 | 10:00 |
| **Order 4** | 1:45 | 3:30 | 5:45 | 7:45 | 9:15 |
| **Order 5** | 2:30 | 5:00 | 6:45 | 8:45 | 10:45 |
| **Order 6** | 2:15 | 4:00 | 6:30 | 9:00 | 10:30 |
| **Order 7** | 2:00 | 4:15 | 6:15 | 8:00 | 10:15 |
| **Order 8** | 3:00 | 4:45 | 7:00 | 8:30 | 11:00 |
| **Order 9** | 1:30 | 4:30 | 7:00 | 9:00 | 10:45 |
| **Order 10** | 2:30 | 4:45 | 6:30 | 8:00 | 11:00 |
| **Order 11** | 2:00 | 4:00 | 5:45 | 8:45 | 9:30 |
| **Order 12** | 1:15 | 3:30 | 6:00 | 7:30 | 10:15 |
| **Order 13** | 3:00 | 3:45 | 5:15 | 8:15 | 9:15 |
| **Order 14** | 2:15 | 3:15 | 6:45 | 7:45 | 9:45 |
| **Order 15** | 1:45 | 5:00 | 6:15 | 7:15 | 10:00 |
| **Order 16** | 2:45 | 4:15 | 5:30 | 8:30 | 10:30 |

**Supplementary Figure 7**

Visual representation of probe order conditions assigned to participants with timestamps of each delivered probe. Participants were randomly assigned to a different probe order condition for each movie clip. Probe orders 1-8 share no overlapping probes. Probe orders 9-16 have overlapping probes from the former probe orders, corresponding to their cell colours, to control for ordering effects across participants.  Probes began at 1 minute and 15 seconds until the 11-minute mark (end of the movie clip)
